# The price of defence: toxins, visual signals and oxidative state in an aposematic butterfly

**DOI:** 10.1101/2021.12.08.471400

**Authors:** Jonathan D. Blount, Hannah M. Rowland, Christopher Mitchell, Michael P. Speed, Graeme D. Ruxton, John A. Endler, Lincoln P. Brower

**Affiliations:** Centre for Ecology & Conservation, College of Life & Environmental Sciences, University of Exeter, Penryn Campus, Cornwall TR10 9FE, UK; Max Planck Institute for Chemical Ecology, Hans-Knöll-Straße 8, Jena, 07745, Germany; Institute of Systems, Molecular and Integrative Biology, University of Liverpool, Liverpool L69 7BE, UK; School of Biology, Sir Harold Mitchell Building, Greenside Place, St Andrews, United Kingdom; Centre for Integrative Ecology, School of Life & Environmental Sciences, Deakin University, Waurn Ponds VIC 3216, Australia; Department of Biology, Sweet Briar College, Sweet Briar VA 24595, USA

**Keywords:** aposematism, cardenolides, cardiac glycosides, resource competition, oxidative lipid damage, *Danaus plexippus*, monarch butterfly, honest signalling

## Abstract

In a variety of aposematic species, the conspicuousness of an individual’s warning signal and the quantity of its chemical defence are positively correlated. This apparent honest signalling is predicted by resource competition models which assume that the production and maintenance of aposematic defences compete for access to antioxidant molecules that have dual functions as pigments and in protecting against oxidative damage. We raised monarch butterflies (*Danaus plexippus*) on their milkweed host-plants (Apocynaceae) with increasing quantities of cardenolides to test whether (1) the sequestration of secondary defences is associated with costs in the form of oxidative lipid damage and reduced antioxidant defences; and (2) that reduced oxidative state can decrease the capacity of individuals to produce aposematic displays. In male monarchs conspicuousness was explained by an interaction between oxidative damage and sequestration: males with high levels of oxidative damage become less conspicuous with increased sequestration of cardenolides, whereas those with low oxidative damage become more conspicuous with increased levels of cardenolides. There was no significant effect of oxidative damage or concentration of sequestered cardenolides on female conspicuousness. Our results demonstrate a physiological linkage between the production of coloration and protection from autotoxicity, and differential costs of signalling in monarch butterflies.

## Introduction

Many species of animals, plants and micro-organisms possess chemical defences that reduce the likelihood of their being eaten [1]. Aposematic species, such as the monarch butterfly (*Danaus plexippus*), signal their chemical defences to predators with conspicuous warning colours [2]. In some species, the conspicuousness of the warning signal and the quantity of chemical defence are positively correlated, for example in some dendrobatid frogs [3-5], marine opisthobranchs [6], ladybird beetles [7, 8], and paper wasps *Polistes dominula* [9]. The identification of these relationships has led to renewed interest in the idea that warning signals may inform predators about levels of prey defence and that such signals are honest indicators of defensive capability [10]. Several theoretical studies have, however, predicted the opposite: that well defended prey should reduce investment in signals, because signalling carries an additional conspicuousness cost, and such prey stand a good chance of surviving any attack unharmed [11-13]. Negative signal-defence correlations have been reported across some Dendrobatid poison frogs [14] including *Dendrobates granuliferus* [15]. Using a novel theoretical model, Blount et al [16] could explain both the positive and negative correlations between warning colours and chemical defences if these two traits are linked through the competitive use of a shared resource. One possible shared resource is energy, which can be limiting for the sequestration or biosynthesis of toxins [17]. Alternatively, sequestration of toxins could impose metabolic costs in terms of oxidative lipid damage and reduced antioxidant defences – an idea that has received limited empirical attention in regard to warning signals.

Many plant allelochemicals are powerful pro-oxidants, which, when ingested, can cause oxidative lipid damage [18]. In their resource competition framework, Blount et al [16] envisaged that pigments used in prey warning signals (e.g., carotenoids, flavonoids, melanins, and pteridines) might play a dual role imparting colour as well as acting as antioxidants that prevent self-damage when storing toxins. In their model, if antioxidants are required to enable high levels of toxicity, signal reliability can be explained if the brightest and most toxic species gain access to more of the limiting resource than those that are less bright and less toxic (figure 1a, c). The prediction of signal honesty breaks down in their model when prey have very abundant resources, in which case it pays to divert these resources increasingly into toxins and not into warning signals, because less conspicuous but highly toxic prey encounter predators less often and have higher chances of surviving attacks (figure 1b, d [16]). Directly controlling antioxidant availability is challenging experimentally, consequently it is unclear whether or not colour-toxin relationships are influenced by the resource state of prey [14, 15], and has not yet been examined experimentally [11, 12, 16, 17, 19].

**Figure 1.**
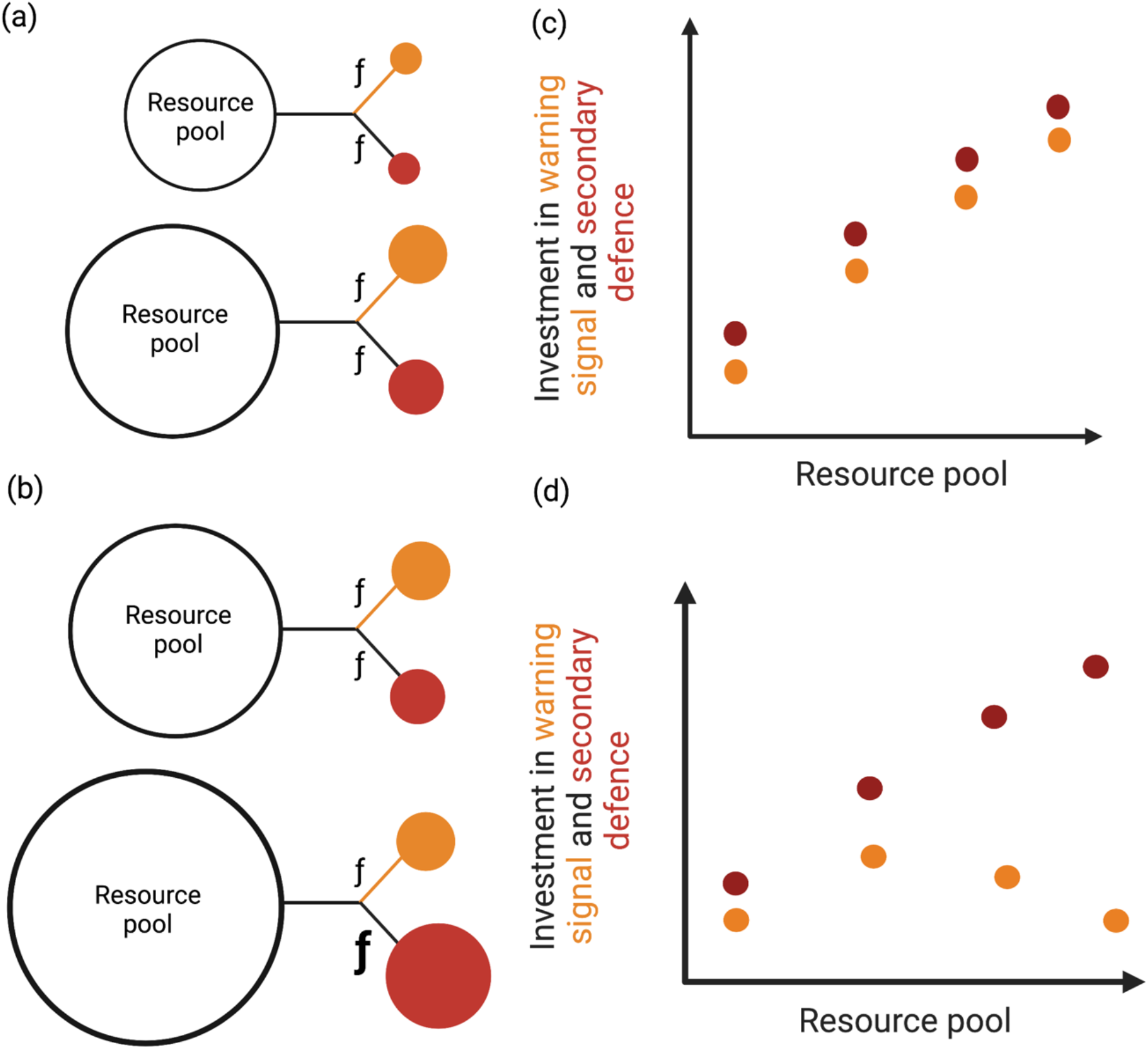
The resource allocation trade off model by Blount et al (2009) predicts that individual prey acquire antioxidant resources from their environment (represented by the circle resource pool), which they divide via genetically or physiologically controlled flow rates (*f*) between sequestering defensive toxins (red circle) and in using antioxidant to producing pigments used for warning signals (orange circle). Each individual or population has access to a given level of resources (represented by the size of the circle) that they allocate to each defence type. In scenario (a) prey allocate resources equally to colour and toxicity. Those with more resources are able to allocate more resources than prey with a smaller resource pool which results in (c) a positive correlation between colour and toxicity, with only the most resource rich prey having the most colourful and toxic defences. In scenario (b) resource availability is very high for some prey (represented by the largest circle resource pool) and, in this situation, those prey with excess resources invest more in toxins than warning signals (represented by the larger *f* and red circle), and results in (d) a negative correlation between colour and toxicity. Blount et al (2009) suggest that this is because highly toxic prey can protect themselves sufficiently even with low levels of warning colours.

We manipulated diet-derived toxins available to an individual to test whether toxin sequestration and the production and maintenance of warning signals are linked via oxidative state. We achieved this by using a model aposematic system, the monarch butterfly (*Danaus plexippus*), that selectively metabolises cardenolides present in its host-plants and stores them in its body [20, 21]. We used the known inter- and intraspecific variability in cardenolide concentration and cardenolide polarity of the monarch’s larval milkweed host-plants (Asclepiadaceae; [21, 22]) to manipulate the toxins available for an individual to sequester. We test whether (i) the quantity of sequestered secondary defence by caterpillars is associated with levels of oxidative lipid damage and antioxidant defences in adults; and (ii) whether oxidative lipid damage incurred during sequestration affects the capacity of adult butterflies to produce aposematic displays.

Because there is substantial genetic variation in sequestration ability, and because evolutionary history and contemporary species interactions may influence patterns of cardenolide sequestration [23] we used monarchs from a single population. We reared them on four milkweed species (see table 1) that show within- and between-species, and among-population variation in cardenolide concentration, diversity, and polarity [22, 24-26]. Based on the published records of biological activity and chromatic quantification (e.g., HPLC and thin layer chromatography) of cardenolides of milkweeds in different populations/latitudes [27, 28], our four plant species were used to represent a gradient of low to high cardenolide defences, with the aim to create different costs of sequestration. We predicted that concentrations of cardenolides in the body and wings of newly eclosed adults would differ according to the cardenolides of the host-plant on which they were reared (see methods): *A. curassavica* (*Ac*) ≥. *A. syriaca* (*As*) > *A. incarnata* (*Ai*) ≥. *A. tuberosa* (*At*).

**Table 1.** The four species of Asclepias used in thh rearing experiments with descriptions of their use by monarchs and phytochemistry.

| Asclepias species | Description and cardenolide content |
| --- | --- |
| *Asclepias curassavica* | A critical hostplant worldwide and contains a diversity of cardenolides, including apolar voruscharin which is detrimental to monarch performance [21, 59-62] |
| *A. syriaca* (common milkweed) | Principal native food plant of the monarch east of the Rocky Mountains and is characterized by more polar cardenolides though also contains the apolar labriformin [63, 64] |
| *A. incarnata* (swamp milkweed) | Preferred hostplant in the Midwest USA and has low concentrations of cardenolides [58]; |
| *A. tuberosa* (butterfly milkweed) | An occasional native foodplant in the wild with low concentrations of cardenolides [21, 65] |

Because sequestration, modification, and storage of allelochemicals are believed to be oxidatively stressful for many chemically defended organisms [18], we predicted that body levels of oxidative lipid peroxidation would differ amongst individuals according to host-plant, as follows: *Ac*≥.*As*>*Ai*≥.*At*. In turn, we predicted that survival to eclosion would differ according to host-plant as follows: *At*≥.*Ai*>*As*≥.*Ac*, and that the redness, luminance, and conspicuousness of the warning signals would differ: *Ac*≥.*As*>*Ai*≥.*At*. In addition to testing for treatment effects, we examined associations between individual levels of cardiac glycosides (CGs), oxidative state, and warning signal redness, luminance, and conspicuousness. We predicted a positive association between levels of CGs and lipid peroxidation, a negative association of CGs with antioxidant defences, and that highly toxic prey will be the most conspicuous [16].

## Materials and methods

### (a) Milkweed plants

Milkweeds (*A. incarnata, A. tuberosa, A. curassavica, and A. syriaca*) were locally sourced in Amherst, Virginia, USA. We did not measure cardenolide content in the leaves of the plants used in the experiment, but used published records about the spatial repeatability of cardenolide defences that take into account latitudinal clines [27-29]. From these records we predicted that for milkweeds grown in Mid-Atlantic and Southeastern regions of the United States, total cardenolide content would vary in the order *A. curassavica* (*Ac*) ≥. *A. syriaca* (*As*) > *A. incarnata* (*Ai*) ≥. *A. tuberosa* (*At*). The plants were grown without the use of pesticides as described previously [30]. All plants were healthy with undamaged leaves at the start of the experiment.

### (b) Milkweed Feeding Assay and preparation of samples

Eggs were obtained from wild caught Monarch females in Virginia (USA) that were kept in captivity until they laid eggs following previous methods [31]. Larvae were reared by LPB on single plants under the same conditions. All larvae had abundant food during the experiment. Larvae were monitored daily for pupation starting at Day 12.

Upon eclosion, the date and the individual’s sex were recorded. Adults were placed in glassine envelopes and frozen on dry ice until dead, then stored in a freezer at -80°C before being shipped to the UK on dry ice. Adults were stored at -80°C in JDB’s laboratory until they were analysed.

For analysis, the gut was removed, and the head, thorax and abdomen (hereafter, ‘body’) were bisected longitudinally. One half of the body was returned to storage at -80°C for later analysis of cardenolides. The other half of the body was immediately homogenised in ice-cold phosphate-buffered saline (PBS; 5 % w/v), for analyses of oxidative stress. The wings were removed using dissecting scissors and shipped on dry ice to HMR’s laboratory for analyses of colouration (see below). Wings were then returned (on dry ice) to JDB’s laboratory for cardenolide analysis.

### (c) Determination of oxidative stress and cardenolide concentration

Determining oxidative state requires a range of assays including antioxidant defences and oxidative damage. We assayed three such components: total superoxide dismutase (SOD), which are metalloenzymes that catalyse the dismutation of superoxide into oxygen and hydrogen peroxide and form a crucial part of intracellular antioxidant defences; total antioxidant capacity (TAC), which measures the activity of low molecular weight chain-breaking antioxidants including ascorbate, a-tocopherol, carotenoids and flavonoids, and therefore gives an overview of the antioxidant status of individuals (including diet-derived antioxidants); and malondialdehyde (MDA), which is formed by the β-scission of peroxidised polyunsaturated fatty acids, and therefore is a definitive marker of oxidative damage.

#### (i) Total superoxide dismutase (SOD)

Total SOD was assayed by measuring the dismutation of superoxide radicals generated by xanthine oxidase and hypoxanthine. We followed the instructions of the kit (#706002; Cayman Chemical, Michigan, USA), except that tissue homogenates were further diluted with the supplied sample buffer (1:100 v/v) to ensure that SOD activity fell within the range of the standard curve. Samples were assayed in duplicate and are reported as units of SOD activity (U) per mg tissue.

#### (ii) Total antioxidant capacity (TAC)

TAC was assayed by measuring the capacity of tissue homogenate to inhibit the oxidation of 2,2’-azino-di-[3-ethylbenzthiazoline sulphonate] (#709001; Cayman Chemical). The homogenate was further diluted with sample buffer (1:5 v/v) to bring the absorbance values within the range of the standard curve. Samples were assayed in duplicate, as per the kit instructions. Data are reported as nmols of TAC activity (Trolox equivalents) per g tissue.

#### (iii) Malondialdehyde (MDA)

MDA was measured using HPLC with fluorescence detection as described previously for tissue homogenates (e.g. [32]). All chemicals were HPLC grade, and chemical solutions were prepared using ultra pure water (Milli-Q Synthesis; Millipore, Watford, UK). Samples were derived in 2 mL capacity polypropylene screw-top microcentrifuge tubes. For tissue homogenates and for the standard (1,1,3,3-tetraethoxypropane, TEP; see below) a 20 μl aliquot was added to 20 μl butylated hydroxytoluene solution (BHT; 0.05% w/v in 95 % ethanol), 160 μl phosphoric acid solution (H_3_PO_4_; 0.44 *M*), and 40 μl thiobarbituric acid solution (TBA; 42 m*M*). Samples were capped, vortex mixed for 5 s, then heated at 100 °C for 1 h in a dry bath incubator to allow formation of MDA-TBA adducts. Samples were then cooled on ice for 5 min, before 80 μl *n*-butanol was added and tubes were vortex mixed for 20s. Tubes were centrifuged at 12,000 x g and 4 °C for 3 min, before the upper (*n*-butanol) phase was collected and transferred to an HPLC vial for analysis. Samples (10 μl) were injected into a Dionex HPLC system (Dionex Corporation, California, USA) fitted with a 5μm ODS guard column and a Hewlett-Packard Hypersil 5μ ODS 100 × 4.6 mm column maintained at 37 °C. The mobile phase was methanol-buffer (40:60, v/v), the buffer being a 50 m*M* anhydrous solution of potassium monobasic phosphate at pH 6.8 (adjusted using 5 *M* potassium hydroxide solution), running isocratically over 3.5 min at a flow rate of 1 ml min^-1^. Data were collected using a fluorescence detector (RF2000; Dionex) set at 515 nm (excitation) and 553 nm (emission). For calibration a standard curve was prepared using a TEP stock solution (5 m*M* in 40 % ethanol) serially diluted using 40 % ethanol. Data are presented as nmols MDA per g tissue.

#### (iv) Cardenolide analysis

For cardenolide analysis, samples (individual wings and the remaining half of the body) were dried in an oven at 60 °C for 16 h. The dried material was ground to a fine powder using a pestle and mortar, and weighed to the nearest 0.0001 g using a GR-202 electronic balance (A&D Instruments Ltd., Abington, UK). The samples were placed in screw-top polypropylene tubes for de-fatting and cardenolide extraction. To remove fats, 2 ml hexane was added and samples were incubated at 35°C for 30 min, before being centrifuged at 10,000 x g and 4ºC for 5 min; the supernatant was collected and discarded. Ethanol (1.9 ml) containing 20 μg digitoxin (#D5878; Sigma-Aldrich, Dorset, UK) as an internal standard were added to the pellet and it was briefly vortexed to extract cardenolides (Ritland 1994). Samples were placed in an XUB5 ultrasonic bath (Grant Instruments, Shepreth, UK) at 60°C for 60 min, followed by further centrifugation (9,600 x g and 4ºC for 5 min). The supernatant was removed and evaporated to dryness using a Savant ISS 110 SpeedVac Concentrator at room temperature (Thermo Scientific, Altrincham, UK). Samples were then dissolved in 0.5 ml methanol by vortexing, and 50 μl of the solution was injected into a Dionex HPLC system fitted with a Waters Spherisorb 5μm ODS2 column (150 × 4.6 mm) maintained at 20°C. A multistep gradient of acetonitrile (ACN) and ultra pure water was used as the mobile phase, as follows: initial concentration 20 % ACN held for 5 minutes, ramp to 70 % ACN at 20 mins and held until 25 minutes, ramp to 95 % ACN at 30 mins and held until 35 mins, returning to 20 % ACN at 40 mins, with an end time of 50 mins. The flow rate was 0.7 ml min^-1^. Peaks were collected using a PDA-100 photodiode array detector (Dionex) at an absorbance value of 218 nm. Spectral data between 200-400 nm were collected, and cardenolides were identified as peaks with a symmetrical absorbance band with a maximum absorbance between 217-222 nm [33, 34]. Data are presented as μg total cardenolides per 0.1 g tissue.

### (d) Measurements and visual modelling of colouration

The reflectance of the forewings of each butterfly, and of four leaves from *Asclepias syriaca* (as a representative background that the butterflies can rest on. Obtained from A. Agrawal), were measured using an Ocean Optics USB2000 spectrophotometer, with specimens illuminated at 45º to normal by a DH1000 balanced halogen deuterium light source. The measuring spot diameter was 3 mm, with spectra recorded at 0.34nm intervals from 300 to 700nm and measured relative to a WS-1 reflectance standard. For each wing, 3-4 non-overlapping measurements were taken per orange segment of the wing (figure s1; total number of measurement spots per forewing = 13-14). For the leaves we took eleven non-overlapping measurements. Spectrophotometry data were recorded using Ocean Optics OOBase32.

To determine differences in chroma between treatments, we modelled the predicted photon catches of a generalized passerine bird (passerines being the main avian predators of monarchs [35]) for each spectrum following the Vorobyev-Osorio model [36], using the R package PAVO [37]. We used the cone types of a blue tit, *Cyanistes caeruleus* [38], longwave, LWS, λmax 563nm; mediumwave, MWS, λmax 503 nm; shortwave, SWS, λmax 448 nm; ultraviolet, UVS, λmax 371 nm; and double dorsal, DD λmax 563 nm]. We calculated redness as the ratio of long wave to medium wave cone responses. We also calculated a measure of chromatic conspicuousness as ΔS between the mean colour of the forewings of the butterflies and mean colour of the *A. syriaca* leaves using the Vorobyev-Osorio colour discrimination model, which is based on evidence that colour discrimination is determined by noise arising in the photoreceptors and is independent of light intensity. We used a Weber fraction value of 0.05 for the most abundant cone type [39]. We calculated luminance as the response of the dorsal double cone [40].

### (e) Data analyses

Left- and right-wing cardenolide concentration (from here on CG) were significantly positively correlated (r = 0.96, t = 20.75, df = 38, p = 2.2e-16). There was no significant correlation between asymmetry in wing CG concentrations and the mean CG of the wings (r = 0.10, t = 0.64, df = 38, P = 0.52), and so for each individual we calculated a mean wing CG concentration. Left- and right-wing redness were significantly positive correlated (r = 0.85, t =10.27, df = 38, P < 0.0001), as were left- and right-wing conspicuousness (r = 0.86, t = 0.33, df = 38, P < 0.0001), and left- and right-wing luminance (r = 0.47, t = 3.32, df = 38, P = 0.002). For each individual we calculated a mean forewing redness, conspicuousness, and luminance.

We log transformed malondialdehyde (MDA) and wing and body CG to normalise the distribution of the residuals. We analysed the effect of foodplant on total CG content, bodyCG and wingCG content using a GLM with a Gaussian distribution and identity link function. We set *A. tuberosa* as the level with which we compared the other plants, because of its low cardenolide content. Eclosion success or failure was analysed with a binomial GLM. To analyse how cardenolides and oxidative stress affected colour metrics in the butterflies we constructed a full model that included all pairwise interactions and a three-way interaction between MDA x total CG concentration x sex and compared this to a simple model that included only the fixed effects of MDA, total CG concentration and sex. We compared the reduced model to the full model by using information criteria (AIC; see supplementary tables s1-s13). We visualized the interaction between two continuous variables using the rsm package in the R [41]. All data were analysed using R version 4.0.4.

## Results

### CG concentrations

In support of our first hypothesis, concentrations of cardenolides in the body and wings of newly eclosed adults differed according to which host-plant species they were reared on (Figure 2a; F_(4,36)_ = 7.42, P = 0.0002). Monarchs reared on *Ac* and *As* sequestered significantly more cardenolides in total than those reared on *At* (*Ac vs At:* estimate = 822.9 ± 184.3, t = 4.47, P = 7.6e-05; *As* vs *At*: estimate = 726.6 ± 187.1, t = 3.88, P =0.0004). Those reared on *Ai* did not sequester significantly different total amounts of CGs to those on *At* (estimate = 213.9 ± 189.2, t = 1.13, P = 0.27). Males tended to sequester more cardenolides in total than females, but this was not significant at the alpha 0.05 level (estimate = 243.5 ± 133.4, t = 1.825, P = 0.076).

**Figure 2.**
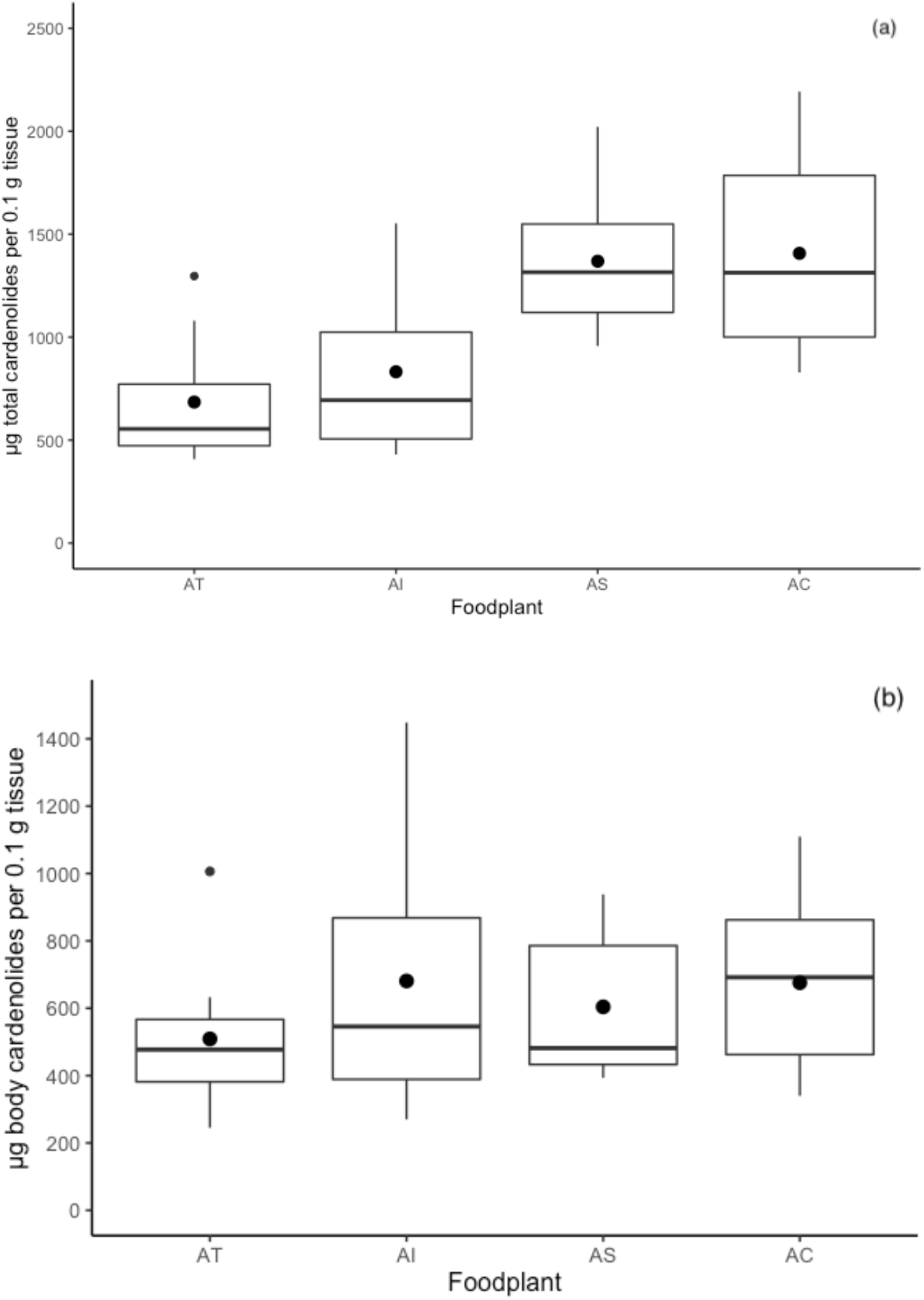
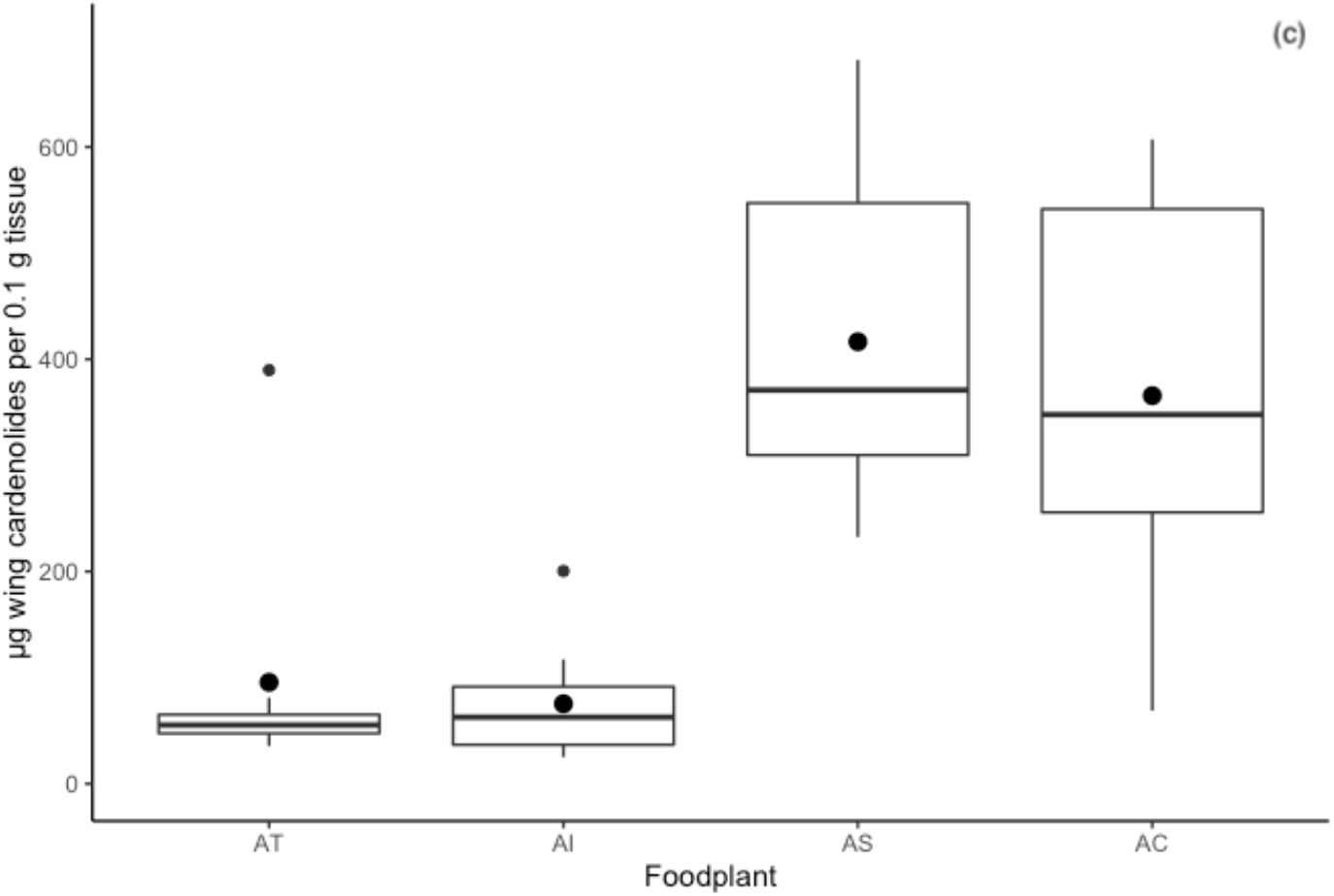
cardenolide sequestration by monarchs feeding on *Ascelpias tuberosa* (AT), *A. Incarnata* (AI), *A. Syriaca* (AS), and *A. curassavica* (AC); a) total cardenolide (μg total cardenolides per 0.1 g tissue) measured in the wings and bodies, b) body cardenolide content, c) wing cardenolide content. Boxplots show the median, interquartile range, and the whiskers represent the largest and smallest value within 1.5 times the 25^th^ and 75^th^ percentile. Outliers are represented by small black dots. The mean cardenolide concentration is represented by the larger black dot.

Body CG concentration tended to differ between food plants, but this was not significant at the alpha 0.05 level (Figure 2b; F_(4,37)_= 2.471, P = 0.061). Wing CG concentration significantly varied between host-plants (Figure 1c; F_(4,36)_= 19.25, P <0.001; Fig. 2c). Wing CGs were significantly higher in *A*c and *As* compared to At (*Ac* vs *At*: estimate = 1.59 ± 0.29, t = 5.48, P < 0.0001; *As* vs *At*: estimate = 1.78 ± 0.30, t = 6.05, P = <0.001), but did not differ significantly between *At* and *Ai* (estimate = -0.05 ± 0.29, τ = −0.18, p=0.38).

### Survival to pupation and eclosion

We predicted that sequestration of cardenolides would impose a cost that would be associated with reduced survivorship to eclosion. Successful eclosion was significantly higher in monarchs reared on *At* (89%) compared with those reared on *As* (20%; estimate = -2.06 ± 0.751, t = 2.75, P = 0.009), while there was no significant difference between *Ai* (70%) and *At* (89%, estimate = -0.696 ± 0.728, t = -0.956, P = 0.34) or *At* (89%) and *Ac* (62%; estimate = -0.927 ± 0.690, t = -1.344, P = 0.19). Survival to pupation did not differ significantly according to host-plant species (0% *At*, 23% *Ai*, 33% *As*, 27% *Ac*; Fisher’s exact test P = 0.28).

### CG and oxidative stress

#### Malondialdehyde (MDA)

Contrary to our prediction that oxidative lipid damage would differ by hostplant, we found no significant effect of hostplant or sex on MDA (main effect F_4, 37_ = 0.77, P = 0.55). In line with our prediction of a positive association between concentration of sequestered CGs and markers of oxidative damage, across all treatment groups individuals that sequestered higher total concentrations of CGs had higher levels of MDA (figure 3; main effect: F_2, 38_ = 4.08, P = 0.025; CG: estimate = 0.0005 ± 0.0002, t = 2.51, P = 0.016). There was no significant difference between the sexes (estimate = -0.3097 ± 0.198, t = -1.57, P = 0.13).

**Figure 3.**
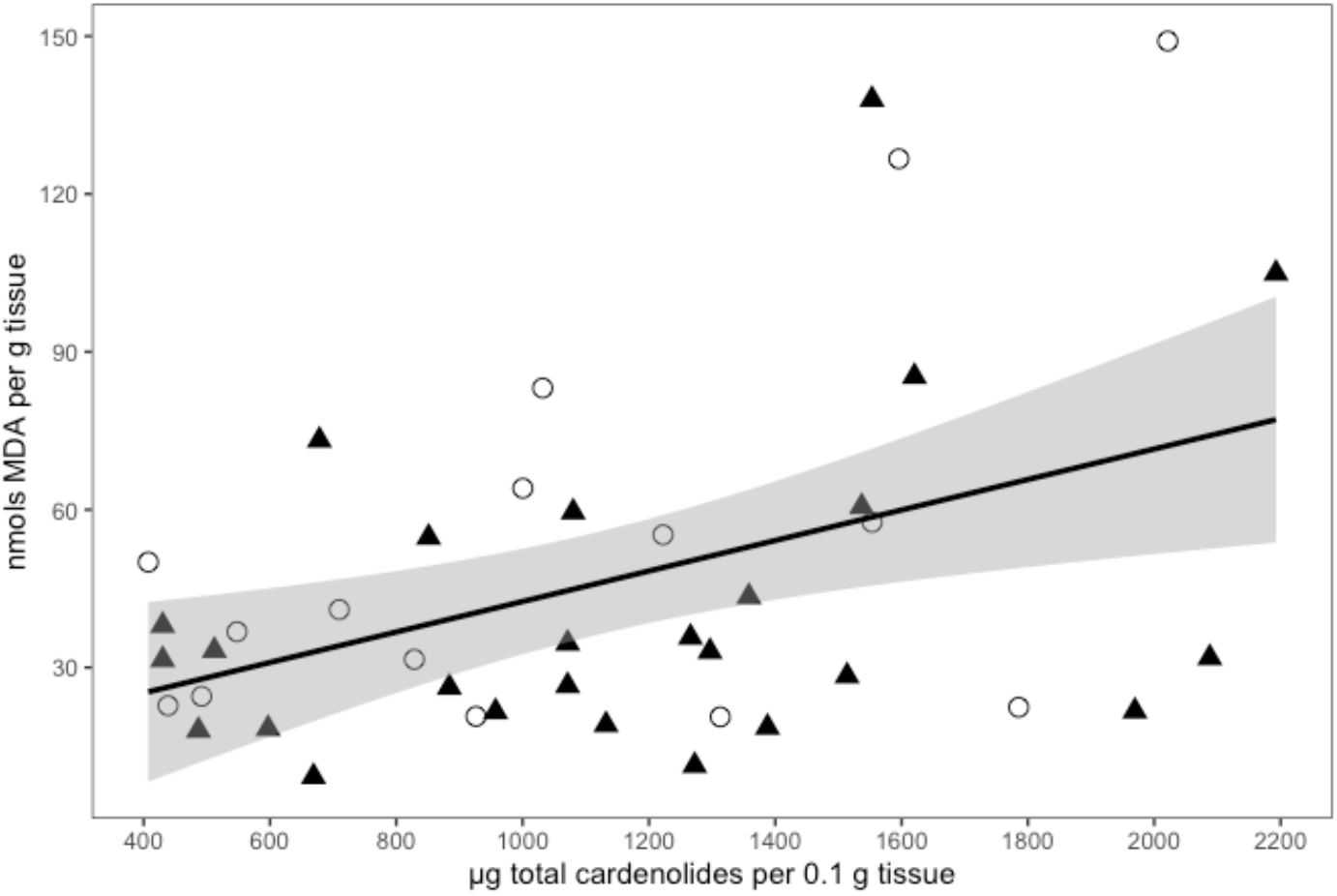
Total cardenolide concentration for males (filled triangle) and females (open circle) and Malondialdehyde (MDA) concentration. Line is the smoothed conditional mean with 95% confidence intervals.

#### Superoxide dismutase activity (SOD)

There was no significant effect of foodplant or sex on SOD activity (main effect F_4, 37_ = 0.28, P = 0.89). We found no significant effect of total CG concentration or sex on SOD activity (main effect: F_2,38_ = 0.52, P = 0.60).

#### Total antioxidant capacity (TAC)

There was no significant effect of foodplant or sex on TAC (: F_(4,37)_= 0.42, P = 0.79) or total CG concentration (F_2,38_ = 0.19, P = 0.83).

#### Sequestration, oxidative stress and warning signals

Because malondialdehyde was the only marker associated with individual levels of sequestration, we proceeded to analyse its association with warning signals, but did not conduct analyses on superoxide dismutase activity or total antioxidant capacity effects on warning signals. Monarchs varied significantly in conspicuousness (chroma (ΔS): F_3,35_ =17.85 P < 0.0001). Males were significantly more conspicuous in chroma (ΔS) than females (estimate = 5.30 ± 0.82, t = 6.49, P < 0.001), and both sexes decreased in conspicuousness with increasing MDA, though this was not significant at the alpha 0.05 level (estimate = -1.26 ± 0.65, t = -1.95, p = 0.06). There was no significant effect of CG concentration on wing conspicuousness (estimate = -0.00002 ± 0.0008, t = -0.03, P = 0.98). Because of the significant sexual dimorphism in colour and sequestration, we separated the analysis by sex and found a significant two way-interaction between MDA and total CG concentration on the conspicuousness (chroma ΔS) of male forewings (main effect: F_3,21_ = 8.54, P = 0.0007; MDA x CG: estimate = -0.003 ± 0.001, t = -3.24, P = 0.004) that is represented by a curved surface (Figure 4). Males with the highest levels of sequestered cardenolides had the most conspicuous warning signals when oxidative damage was lowest, and reduced their warning signal conspicuousness as oxidative damage increased, while those that sequestered the lowest levels of cardenolides showed no change in warning signal conspicuousness across all levels of oxidative damage. The conspicuousness of males warning signals decreased with increasing sequestration in males with high oxidative damage, whereas conspicuousness increased in males with low oxidative damage with increasing sequestration. There was no significant effect of oxidative damage or concentration of sequestered cardenolides on female forewing conspicuousness (F = _3,10_ = 0.06, P = 0.9819).

**Figure 4.**
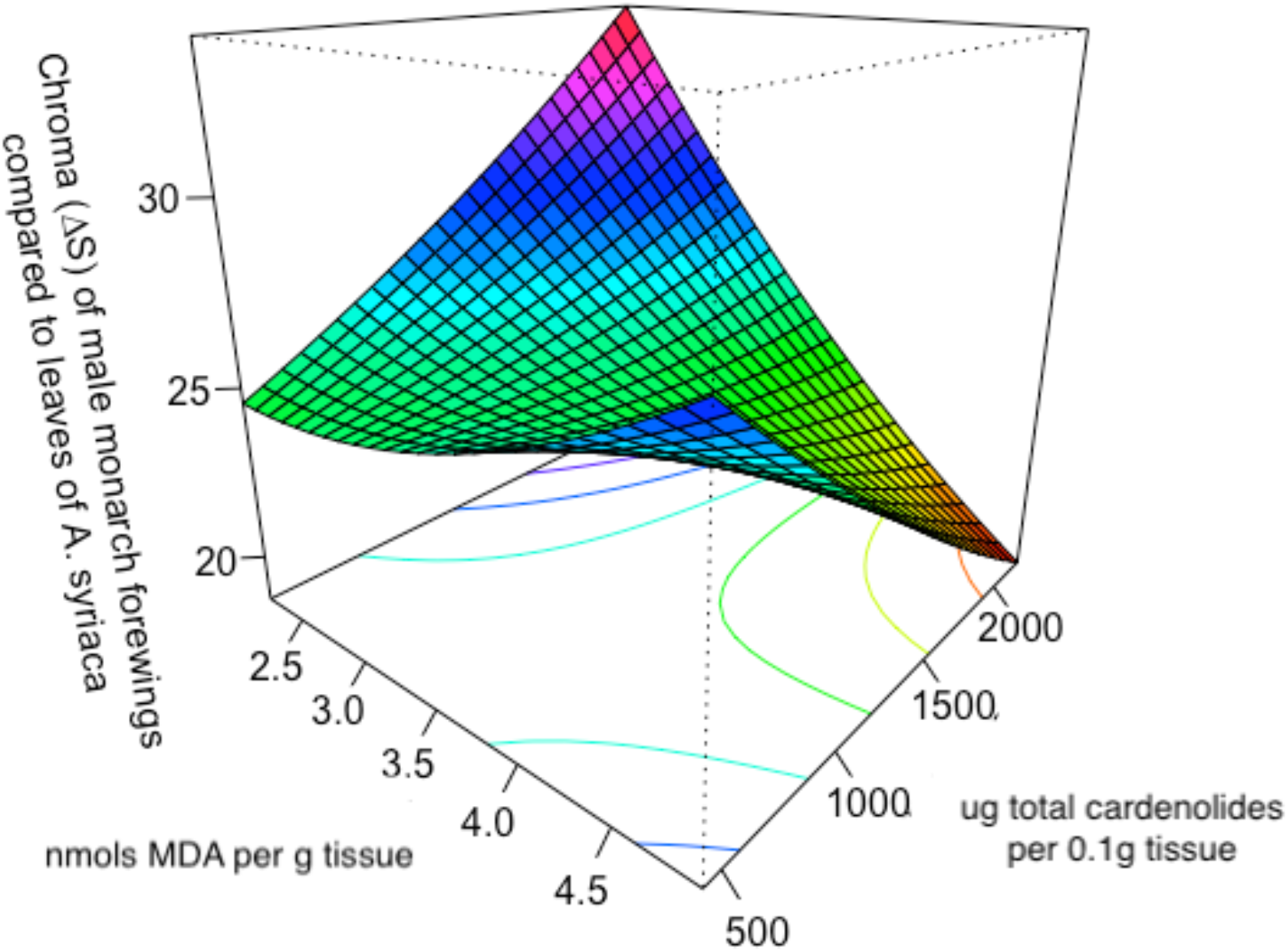
Three-dimensional surface (perspective) plot for the response surface predicting the relationship between the conspicuousness of the male monarch’s forewing, oxidative damage measured as malondialdehyde (MDA), and total cardenolide concentration.

Males also had significantly redder forewings than females (main effect: F_3,35_ = 5.81, P = 0.002; sex estimate = 0.24 ± 0.06, t = 3.82, P = 0.0005), but there was no significant effect of MDA or CG concentration on male and female redness (MDA: estimate = -0.04 ± 0.049, t = -0.86, p = 0.40; CG concentration: estimate = 0.00001 ± 0.00006, t = 0.16, P = 0.87). We found a significant difference in brightness of the monarchs (F_3, 35_ = 10.19, P = 0.00006). Males were significantly brighter than females (estimate = 0.03 ± 0.007, t = 4.37, p = 0.0001) and both sexes decreased in brightness with increasing MDA (estimate = -0.01 ± 0.005, t = - 2.40, p = 0.02). There was no significant effect of CG concentration on wing brightness (estimate = 0.000003 ± 0.000007, t = 0.37, P = 0.70).

## Discussion

We reared individual monarch caterpillars on whole plants from single plant populations of four milkweed species with varying phytochemistry (*Asclepias tuberosa, A. incarnata, A*.*syriaca*, and *A*.*curassavica*; [42]). Monarchs that sequestered higher concentrations of cardenolides experienced higher levels of oxidative damage than those that sequestered lower concentrations (measured as the concentration of malondialdehyde; MDA). Although there is some evidence that cardenolides can be a burden for monarch caterpillars [22, 43], our results are some of the first to show a potential physiological mechanism of oxidative damage as a cost of sequestration for monarchs (see also [21, 29]). We found a trade-off between sequestration and conspicuousness in male monarchs mediated by oxidative damage. Males with high oxidative damage had decreasing conspicuousness with increasing sequestration, whereas males with low oxidative damage showed increased conspicuousness with increasing sequestration. This is the first evidence to support Blount et al’s [16] resource allocation trade-off model – prey with lower oxidative damage are able to allocate more resources to colour and toxicity than prey with higher oxidative damage – and shows that an individual’s ability to deal with oxidative damage can result in colour-toxin correlations in monarch butterflies.

Some authors [44] have rejected the idea that there can be a mechanistic link between warning signals and chemical defences that involve intrinsic costs of production or maintenance of defences because there was no clear mediator. However, condition-dependent expression of colourful signals can arise when the underlying physiological pathways that produce colouration are dependent on the same core cellular processes [16, 45]. We found stable levels of antioxidant defences (superoxidase dismutase and total antioxidant capacity) but increased oxidative damage was associated with reduced warning signal conspicuousness, which suggests that the monarchs that are better able to control pro-oxidant levels can sequester more cardenolides and still invest in warning signals [46]. Similar patterns are observed in *Aristolochia*-feeding pipevine swallowtails butterflies (*Battus philenor*) that store nitrophenanthrenes and have significantly higher tissue levels of carotenoids than related species which mimic them in coloration but do not store toxins [47]. Similarly, the rank order of body concentrations of carotenoids in three species of lepidoptera matches the rank order of dietary exposure to the prooxidant toxin furanocoumarin (8-MOP) [Papilio polyxenes > Spodoptera eridania > Trichoplusia ni; [18]]. Antioxidants are involved in detoxification processes [48] as well as pigment synthesis pathways directly [49], or as cofactors of enzymes [50]. Our results allow us to speculate that antioxidant availability has a role in the biochemistry underlying the variations in warning colouration and toxicity in aposematic animals. However, warning colours are usually regulated by more than one mechanism, and we suggest that this area of research warrants further biochemical study.

It is likely that oxidative state depends on the combination of genetic, environmental and gene by environment (GxE) interactions that determine an individual’s condition. Monarchs show patterns of local adaptation to their hostplants (based on larval growth rate [29]), and also show GxE interactions in sequestration ability [23] which may reflect either a lack of evolutionary history with different species of *Asclepias*, or a physiological trade-off in sequestration ability. Our results suggest that populations of monarchs that are sympatric with high cardenolide milkweeds could be less subject to oxidative damage, either because they have evolved higher antioxidant efficiency, or because they are able to accumulate more antioxidants to provide enhanced antioxidant protection. Further research on this topic could have interesting implications for understanding host shifts and range expansions as well as our understanding of automimicry (the occurrence of palatable ‘cheaters’ in a chemically defended population). Whether automimicry in monarchs arises directly from a deficiency in an individual’s capacity to cope with reactive oxygen species warrants further study [51]. The resource competition model does not investigate or predict sex differences in resource limitation. However, Blount et al [52] found that ladybird females (the larger sex) are more susceptible to resource limitation than males, and hence more likely to signal honestly. In this study we found costs of signalling in male monarchs. Males are slightly larger than females and are the more active sex. Males also sequestered more cardenolides than females. This behavioural difference between the sexes may increase their probability of detection and risk of predation, and also increase their sensitivity to oxidative damage. If predators are sensitive to the combined qualities of toxins and displays when they attack prey, this could mean they avoid highly conspicuous prey and reject highly defended but less conspicuous prey, and instead predate moderately defended and moderately conspicuous individuals, this could maintain the signal variability we observe in males and in other aposematic prey.

Our results depend, to some extent, on the measure of signal conspicuousness measured against the leaves of one of the four hostplants (*A. syriaca*). The differences in conspicuousness of the forewings to the hostplant leaves range from (ΔS) 14 to 29. These ‘suprathreshold’ colour differences are often assumed to scale linearly with colour distance, but this is not the case in some visual systems [53]. Future research could calculate conspicuousness against a range of hostplant species and plant parts, as well as considering predator visual acuity and the effects of increasing viewing distance on warning signal conspicuousness. Experiments testing predator responses to the colour differences and the consequences for prey survival will also be important for determining whether the levels of measured conspicuousness are also perceived as visually distinct at a distance. The relationship between visual signal strength, chemical defence, and predator response warrants further study.

Although we did not measure cardenolide content in milkweed plants, our results confirm previous research showing that the concentration of cardenolides in monarch butterflies varies depending on host-plant chemistry [54, 55], and follows a standard approach to using the natural between species and population variability in plant toxicity to test for costs of sequestration [21, 27, 42]. However, it is methodologically difficult to separate the costs of sequestration from other potential effects of plant allelochemicals [56]. Future work could focus on comparative phytochemical profiles to understand resource availability, and experiments that control nutritional content while manipulating toxin content [57].

In conclusion, our results support the proposal that oxidative state can be a key physiological mechanism that links warning colours to sequestration costs [16], and that the contribution of pigments to antioxidant activity for maintaining redox homeostasis is important [51]. Our results show that specialist herbivores must balance the benefits of plant secondary compounds for sequestration with the burden that these same compounds impose (though see [42, 58]). Future research should examine the possibly of natural selection for oxidative capacity, and whether aposematism evolves in species that have high antioxidant capacity. These studies could provide new theories for understanding the evolution of aposematism when coloration and toxicity do not coevolve. Documenting the costs associated with using secondary defences in natural systems is important for our understanding of the ecology and evolution of aposematism.

## Supporting information

Blount et al Supplementary material

## Acknowledgements

We thank Prof. Linda Fink for her support in drafting the manuscript following the death of Prof. Lincoln Brower. We thank Alfonso Aceves for help with the PAVO coding for visual modelling, and Prof. Anurag Agrawal for leaves of *Ascelpias syriaca*. JDB was supported by a Royal Society University Research Fellowship. HMR, GDR and MPS were supported by NERC (NE/D010 667/1) during data collection, and HMR was supported by a Junior Research Fellowship from Churchill College, Cambridge and the Max Planck Society during data analysis and manuscript preparation.

## Author contributions

**Jonathan D. Blount** conceptualization, methodology, resources, data curation, writing - original draft, project administration, funding acquisition.

**Hannah M. Rowland** methodology, formal analysis, investigation, data curation, writing - original draft, visualization.

**Christopher Mitchell** investigation, data curation.

**Michael P. Speed** conceptualization, methodology, writing - review & editing.

**Graeme D. Ruxton** conceptualization, methodology, writing - review & editing.

**John A. Endler** conceptualization, writing - review & editing, visualization.

**Lincoln P. Brower** conceptualization, methodology, investigation, resources.

## References

[1] Blum, M.S. 1981 Chemical Defenses of Arthropods. New York, Academic Press.

[2] Ruxton, G.D., Allen, W.L., Sherratt, T.N. & Speed, M.P. 2018 Avoiding Attack. The Evolutionary Ecology of Crypsis, Aposematism, and Mimicry. Oxford, UK., Oxford University Press.

[3] Summers, K. & and Clough, M.E. 2001 The evolution of coloration and toxicity in the poison frog family (Dendrobatidae). Proceedings of the National Academy of Sciences 98, 6227–6232.

[4] Santos, J.C. & Cannatella, D.C. 2011 Phenotypic integration emerges from aposematism and scale in poison frogs. Proceedings of the National Academy of Sciences 108, 6175–6180. (doi:10.1073/pnas.1010952108).

[5] Maan, M.E. & Cummings, M.E. 2009 Sexual dimorphism and directional sexual selection on aposematic signals in a poison frog. Proceedings of the National Academy of Sciences 106, 19072–19077. (doi:10.1073/pnas.0903327106).

[6] Cortesi, F. & Cheney, K.L. 2010 Conspicuousness is correlated with toxicity in marine opisthobranchs. Journal of Evolutionary Biology 23, 1509–1518. (doi:doi:10.1111/j.1420-9101.2010.02018.x).

[7] Bezzerides, A. L., McGraw, K.J., Parker, R.S. & Husseini, J. 2007 Elytra color as a signal of chemical defense in the Asian ladybird beetle Harmonia axyridis. Behavioral Ecology and Sociobiology 61, 1401–1408. (doi:10.1007/s00265-007-0371-9).

[8] Winters, A.E., Stevens, M., Mitchell, C., Blomberg, S.P. & Blount, J.D. 2014 Maternal effects and warning signal honesty in eggs and offspring of an aposematic ladybird beetle. Functional Ecology 28, 1187–1196. (doi:doi:10.1111/1365-2435.12266).

[9] Vidal-Cordero, J.M., Moreno-Rueda, G., López-Orta, A., Marfil-Daza, C., Ros-Santaella, J.L. & Ortiz-Sánchez, F.J. 2012 Brighter-colored paper wasps (Polistes dominula) have larger poison glands. Frontiers in Zoology 9, 20. (doi:10.1186/1742-9994-9-20).

[10] Summers, K., Speed, M.P., Blount, J.D. & Stuckert, A.M.M. 2015 Are aposematic signals honest? A review. Journal of Evolutionary Biology, n/a-n/a. (doi:10.1111/jeb.12676).

[11] Leimar, O., Enquist, M. & Sillentullberg, B. 1986 Evolutionary Stability of Aposematic Coloration and Prey Unprofitability - a Theoretical-Analysis. American Naturalist 128, 469–490.

[12] Speed, M.P. & Ruxton, G.D. 2007 How bright and how nasty: Explaining diversity in warning signal strength. Evolution 61, 623–635.

[13] Ruxton, G.D., Speed, M.P. & Broom, M. 2009 Identifying the ecological conditions that select for intermediate levels of aposematic signalling. Evolutionary Ecology 23, 491–501. (doi:10.1007/s10682-008-9247-3).

[14] Darst, C.R., Cummings, M.E. & Cannatella, D.C. 2006 A mechanism for diversity in warning signals: Conspicuousness versus toxicity in poison frogs. Proc. Natl. Acad. Sci. U. S. A. 103, 5852–5857.

[15] Wang, I.J. 2011 Inversely related aposematic traits: reduced conspicuousness evolves with increased toxicity in a polymorphic poison-dart frog. Evolution 65, 1637–1649. (doi:doi:10.1111/j.1558-5646.2011.01257.x).

[16] Blount, J.D., Speed, M.P., Ruxton, G.D. & Stephens, P.A. 2009 Warning displays may function as honest signals of toxicity871–877 p.

[17] Holloway, G.J., de Jong, P.W., Brakefield, P.M. & de Vos, H. 1991 Chemical defence in ladybird beetles (Coccinellidae). I. Distribution of coccinelline and individual variation in defence in 7-spot ladybirds (Coccinella septempunctata). CHEMOECOLOGY 2, 7–14. (doi:10.1007/bf01240660).

[18] Ahmad, S. 1992 Biochemical defence of pro-oxidant plant allelochemicals by herbivorous insects. Biochemical Systematics and Ecology 20, 269–296.

[19] Holen, O.H. & Svennungsen, T.O. 2012 Aposematism and the Handicap Principle. The American Naturalist 180, 629–641. (doi:10.1086/667890).

[20] Malcolm, S.B. & Brower, L.P. 1989 Evolutionary and Ecological Implications of Cardenolide Sequestration in the Monarch Butterfly. Experientia 45, 284–295.

[21] Agrawal, A.A., Böröczky, K., Haribal, M., Hastings, A.P., White, R.A., Jiang, R.-W. & Duplais, C. 2021 Cardenolides, toxicity, and the costs of sequestration in the coevolutionary interaction between monarchs and milkweeds. Proceedings of the National Academy of Sciences 118, e2024463118. (doi:doi:10.1073/pnas.2024463118).

[22] Tao, L., Berns, A.R. & Hunter, M.D. 2014 Why does a good thing become too much? Interactions between foliar nutrients and toxins determine performance of an insect herbivore. Functional Ecology 28, 190–196. (doi:https://doi.org/10.1111/1365-2435.12163).

[23] Freedman, M.G., Choquette, S.-L., Ramírez, S.R., Strauss, S.Y., Hunter, M.D. & Vannette, R.L. 2021 Population-specific patterns of toxin sequestration in monarch butterflies from around the world. bioRxiv, 2021.2010.2015.464593. (doi:10.1101/2021.10.15.464593).

[24] Brower, L.P. 1969 Ecological chemistry. Scientific American 220, 22–29.

[25] Brower, L.P. 1972 Ecological Chemistry of Palatability Cardiac Glycoside Spectrum in Monarch Butterflies, Danaus-Plexippus, and Asclepias Milkweeds. American Zoologist 12, 712–713.

[26] Züst, T., Petschenka, G., Hastings, A.P. & Agrawal, A.A. 2019 Toxicity of Milkweed Leaves and Latex: Chromatographic Quantification Versus Biological Activity of Cardenolides in 16 Asclepias Species. J Chem Ecol 45, 50–60. (doi:10.1007/s10886-018-1040-3).

[27] Soule, A.J., Decker, L.E. & Hunter, M.D. 2020 Effects of diet and temperature on monarch butterfly wing morphology and flight ability. Journal of Insect Conservation 24, 961–975. (doi:10.1007/s10841-020-00267-7).

[28] Woods, E.C., Hastings, A.P., Turley, N.E., Heard, S.B. & Agrawal, A.A. 2012 Adaptive geographical clines in the growth and defense of a native plant. Ecological Monographs 82, 149–168. (doi:https://doi.org/10.1890/11-1446.1).

[29] Freedman, M.G., Jason, C., Ramírez, S.R. & Strauss, S.Y. 2020 Host plant adaptation during contemporary range expansion in the monarch butterfly. Evolution 74, 377–391. (doi:https://doi.org/10.1111/evo.13914).

[30] Pocius, V.M., Debinski, D.M., Pleasants, J.M., Bidne, K.G., Hellmich, R.L. & Brower, L.P. 2017 Milkweed Matters: Monarch Butterfly (Lepidoptera: Nymphalidae) Survival and Development on Nine Midwestern Milkweed Species. Environ. Entomol. 46, 1098–1105. (doi:10.1093/ee/nvx137).

[31] Zalucki, M.P. & Brower, L.P. 1992 Survival of first instar larvae ofDanaus plexippus (Lepidoptera: Danainae) in relation to cardiac glycoside and latex content ofAsclepias humistrata (Asclepiadaceae). CHEMOECOLOGY 3, 81–93. (doi:10.1007/BF01245886).

[32] Blount, J.D. & Pike, T.W. 2012 Deleterious effects of light exposure on immunity and sexual coloration in birds. Functional Ecology 26, 37–45. (doi:https://doi.org/10.1111/j.1365-2435.2011.01926.x).

[33] Malcolm, S.B. & Zalucki, M.P. 1996 Milkweed latex and cardenolide induction may resolve the lethal plant defence paradox. In Proceedings of the 9th International Symposium on Insect-Plant Relationships (eds. E. Städler, M. Rowell-Rahier & R. Bauer), pp. 193–196. Dordrecht, Springer Netherlands.

[34] Rasmann, S., Agrawal, A.A., Cook, S.C. & Erwin, A.C. 2009 Cardenolides, induced responses, and interactions between above- and belowground herbivores of milkweed (Asclepias spp.). Ecology 90, 2393–2404. (doi:10.1890/08-1895.1).

[35] Fink, L.S., Brower, L.P., Waide, R.B. & Spitzer, P.R. 1983 Overwintering Monarch Butterflies as Food for Insectivorous Birds in Mexico. Biotropica 15, 151–153.

[36] Vorobyev, M. & Osorio, D. 1998 Receptor noise as a determinant of colour thresholds. Proc. R. Soc. Lond. Ser. B-Biol. Sci. 265, 351–358.

[37] Maia, R., Gruson, H., Endler, J.A. & White, T.E. 2019 pavo 2: New tools for the spectral and spatial analysis of colour in r. Methods in Ecology and Evolution 10, 1097–1107. (doi:https://doi.org/10.1111/2041-210X.13174).

[38] Hart, N.S., Partridge, J.C., Cuthill, I.C. & Bennett, A.T.D. 2000 Visual pigments, oil droplets, ocular media and cone photoreceptor distribution in two species of passerine bird: the blue tit (Parus caeruleus L.) and the blackbird (Turdus merula L.). J. Comp. Physiol. A-Sens. Neural Behav. Physiol. 186, 375–387.

[39] Silvasti, S.A., Valkonen, J.K. & Nokelainen, O. 2021 Behavioural thresholds of blue tit colour vision and the effect of background chromatic complexity. Vision Research 182, 46–57. (doi:https://doi.org/10.1016/j.visres.2020.11.013).

[40] Osorio, D. & Vorobyev, M. 2005 Photoreceptor spectral sensitivities in terrestrial animals: adaptations for luminance and colour vision. Proceedings of the Royal Society of London, Series B 272, 1745–1752.

[41] Lenth, R.V. 2009 Response-Surface Methods in R, Using rsm. Journal of Statistical Software 32, 1–17. (doi:10.18637/jss.v032.i07).

[42] Erickson, J.M. 1973 The Utilization of Various <em>Asclepias</em> Species by Larvae of the Monarch Butterfly, <em>Danaus Plexippus</em>. Psyche 80, 028693. (doi:10.1155/1973/28693).

[43] Zalucki, M.P., Brower, L.P. & Alonso, A. 2001 Detrimental effects of latex and cardiac glycosides on survival and growth of first-instar monarch butterfly larvae Danaus plexippus feeding on the sandhill milkweed Asclepias humistrata. Ecological Entomology 26, 212–224.

[44] Guilford, T. & Dawkins, M.S. 1993 Are warning colors handicaps? Evolution 47, 400.

[45] Hill, G.E. 2011 Condition-dependent traits as signals of the functionality of vital cellular processes. Ecol Lett 14, 625–634. (doi:10.1111/j.1461-0248.2011.01622.x).

[46] Costantini, D., Fanfani, A. & Dell’omo, G. 2008 Effects of corticosteroids on oxidative damage and circulating carotenoids in captive adult kestrels (Falco tinnunculus). J Comp Physiol B 178, 829–835. (doi:10.1007/s00360-008-0270-z).

[47] Harborne, J.B. & Williams, C.A. 2001 Anthocyanins and other flavonoids. Natural product reports, 18, 310–333.

[48] Enayati, A.A., Ranson, H. & Hemingway, J. 2005 Insect glutathione transferases and insecticide resistance. Insect Mol Biol 14, 3–8. (doi:10.1111/j.1365-2583.2004.00529.x).

[49] Ito, S. & Prota, G. 1977 A facile one-step synthesis of cysteinyldopas using mushroom tyrosinase. Experientia 33, 1118–1119. (doi:10.1007/bf01946005).

[50] Shamim, G., Ranjan, K.S., Pandey, M.D. & Ramani, R. 2014 Biochemistry and biosynthesis of insect pigments. EJE 111, 149–164.

[51] Costantini, D. & Verhulst, S. 2009 Does high antioxidant capacity indicate low oxidative stress? Functional Ecology 23, 506–509. (doi:https://doi.org/10.1111/j.1365-2435.2009.01546.x).

[52] Blount, J.D., Rowland, H.M., Drijfhout, F.P., Endler, J.A., Inger, R., Sloggett, J.J., Hurst, G.D.D., Hodgson, D.J. & Speed, M.P. 2012 How the ladybird got its spots: effects of resource limitation on the honesty of aposematic signals. Functional Ecology 26, 334–342. (doi:10.1111/j.1365-2435.2012.01961.x).

[53] Santiago, C., Green, N.F., Hamilton, N., Endler, J.A., Osorio, D.C., Marshall, N.J. & Cheney, K.L. 2020 Does conspicuousness scale linearly with colour distance? A test using reef fish. Proceedings of the Royal Society B: Biological Sciences 287, 20201456. (doi:doi:10.1098/rspb.2020.1456).

[54] Jones, P.L., Petschenka, G., Flacht, L. & Agrawal, A.A. 2019 Cardenolide Intake, Sequestration, and Excretion by the Monarch Butterfly along Gradients of Plant Toxicity and Larval Ontogeny. Journal of Chemical Ecology 45, 264–277. (doi:10.1007/s10886-019-01055-7).

[55] Brower, L.P., Nelson, C.J., Seiber, J.N., Fink, L.S. & Bond, C. 1988 Exaptation as an alternative to coevolution in the cardenolide-based chemical defense of monarch butterflies (Danaus plexippus L.) against avian predators. In Chemical mediation of coevolution (ed. K.C. Spencer). London, Harcourt Brace Jovanovich.

[56] Bowers, M.D. 1992 The evolution of unpalatability and the cost of chemical defense in insects. In Insect chemical ecology: an evolutionary approach. Chapman & Hall, New York, New York, USA. (eds. B.D. Roitberg & M.B. Isman), pp. 216–244.

[57] Pokharel, P., Steppuhn, A. & Petschenka, G. 2021 Dietary cardenolides enhance growth and change the direction of the fecundity-longevity trade-off in milkweed bugs (Heteroptera: Lygaeinae). Ecology and evolution 11, 18042–18054. (doi:https://doi.org/10.1002/ece3.8402).

[58] Petschenka, G. & Agrawal, A.A. 2015 Milkweed butterfly resistance to plant toxins is linked to sequestration, not coping with a toxic diet. Proceedings of the Royal Society B: Biological Sciences 282. (doi:10.1098/rspb.2015.1865).

[59] Seiber, J.N., Benson, J.M., Roeske, C.A. & Brower, L.P. 1975 Qualitative and Quantitative Aspects of Milkweed Cardenolide Sequestering by Monarch Butterflies. Abstracts of Papers of the American Chemical Society 170, 103–103.

[60] Roeske, C.N., Seiber, J.N., Brower, L.P. & Moffitt, C.M. 1976 Milkweed Cardenolides and Their Comparative Processing by Monarch Butterflies (Danaus plexippus L.). In Biochemical Interaction Between Plants and Insects (eds. J.W. Wallace & R.L. Mansell), pp. 93–167. Boston, MA, Springer US.

[61] Faldyn, M.J., Hunter, M.D. & Elderd, B.D. 2018 Climate change and an invasive, tropical milkweed: an ecological trap for monarch butterflies. Ecology 99, 1031–1038. (doi:https://doi.org/10.1002/ecy.2198).

[62] Agrawal, A. 2017 Monarchs and Milkweed A Migrating Butterfly, a Poisonous Plant, and Their Remarkable Story of Coevolution, Princeton University Press.

[63] Geest, E.A., Wolfenbarger, L.L. & McCarty, J.P. 2019 Recruitment, survival, and parasitism of monarch butterflies (Danaus plexippus) in milkweed gardens and conservation areas. Journal of Insect Conservation 23, 211–224. (doi:10.1007/s10841-018-0102-8).

[64] Malcolm, S.B., Cockrell, B.J. & Brower, L.P. 1989 Cardenolide Fingerprint of Monarch Butterflies Reared on Common Milkweed, Asclepias-Syriaca-L. Journal of Chemical Ecology 15, 819–853.

[65] Brower, L.P., Brower, J.V.Z. & Corvino, J.M. 1967 Plant poisons in terrestrial food chain. Proc. Natl. Acad. Sci. U. S. A. 57, 893–898.

