## Supplementary material for "The price of defence: toxins, visual signals and oxidative state in an aposematic butterfly": Blount et al Supplementary material

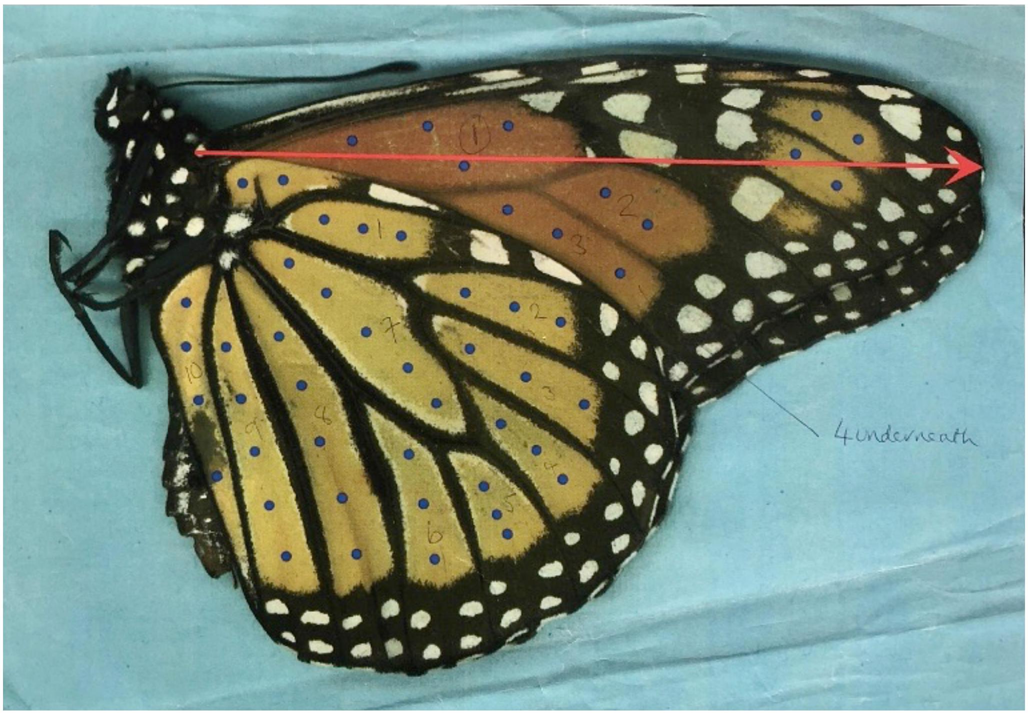


**Figure s1.** The spectrophometric measurement locations on the fore and hindwings of an example fully intact adult monarch butterfly. Isolated butterfly wings were used for spectrophotometry analysis. The blue dots show the locations of measurements within each panel, and the numbers are the different panels within the wing.

.

**Table S1**. the main effects GLM on total CG and foodplant (bold represents minimal adequate model)

| Model | AIC |
|---|---|
| cgconcOverall ~ food + sex + food:sex, | 613.24 |
| **cgconcOverall ~ food + sex** | **612.53** |

**Table S2**. the main effects GLM on body CG and foodplant (bold represents minimal adequate model)

| | |
|---|---|
| logCGbody ~ food + sex + food:sex | 47.8165 |
| logCGbody ~ food + sex | 47.72835 |

**Table S3.** the main effects GLM on wing CG and foodplant (bold represents minimal adequate model)

| | |
|---|---|
| logCGwingAv ~ food + sex + food:sex | 86.54536 |
| **logCGwingAv ~ food + sex** | **83.4386** |

**Table S4.** Binomial GLM on pupation (bold represents minimal adequate model)

| | |
|---|---|
| ok ~ food + sex + food:sex | 59.73983 |
| ok ~ food + sex | 55.6073 |
| **ok ~ food** | 53.82755 |

**Table S5**. the "main effects" GLM on food and MDA (bold represents minimal adequate model)

| | |
|---|---|
| logMDA ~ food + sex + food:sex | 90.03 |
| logMDA ~ food + sex | 90.29 |

**Table S6** the "main effects" GLM on CG and MDA (bold represents minimal adequate model)

| | |
|---|---|
| logMDA ~ sex + cgconcOverall + cgconcOverall:sex | 82.203 |
| **logMDA ~ sex + cgconcOverall** | **80.37** |

**Table S7** the "main effects" GLM on food and SOD (bold represents minimal adequate model)

| | |
|---|---|
| SOD ~ food + sex + food:sex | 178.26 |
| **SOD ~ food + sex** | **173.06** |

**Table S8** the "main effects" GLM on food and SOD (bold represents minimal adequate model)

| | |
|---|---|
| SOD ~ sex + cgconcOverall + cgconcOverall:sex | 167.32 |

| | |
|---|---|
| **SOD ~ sex + cgconcOverall** | **165.8** |

**Table S9.** the "main effects" GLM on food and TAC (bold represents minimal adequate model)

| | |
|---|---|
| TAC ~ food + sex + food:sex | 573.02 |
| **TAC ~ food + sex** | **568.2** |

**Table S10.** the "main effects" GLM on CG and TAC (bold represents minimal adequate model)

| | |
|---|---|
| TAC ~ sex + cgconcOverall +cgconcOverall:sex | 554.47 |
| **TAC ~ sex + cgconcOverall** | **553.36** |

**Table S11.** Oxidative stress and colouration (bold represents minimal adequate model)

| | |
|---|---|
| FW.consp ~ logMDA + cgconcOverall + sex + logMDA:cgconcOverall + logMDA:sex + cgconcOverall:sex+ logMDA:cgconcOverall:sex | 183.7716 |
| FW.consp ~ logMDA + cgconcOverall + sex + logMDA:cgconcOverall + logMDA:sex + cgconcOverall:sex, | 185.1388 |
| FW.consp ~ logMDA + cgconcOverall + sex | 184.3485 |

**Table s12.** GLM MDA and forewing redness (bold represents minimal adequate model)

| | |
|---|---|
| FW.red ~ logMDA + cgconcOverall + sex + logMDA:cgconcOverall + logMDA:sex + cgconcOverall:sex + logMDA:cgconcOverall:sex, | -15.915 |
| FW.red ~ logMDA + cgconcOverall + sex + logMDA:cgconcOverall + logMDA:sex + cgconcOverall:sex, | -16.165 |
| **FW.red ~ logMDA + cgconcOverall + sex** | **-17.345** |

**Table s13**. GLM MDA and forewing brightness (bold represents minimal adequate model)

| | |
|---|---|
| FW.bright ~ logMDA + cgconcOverall + sex + logMDA:cgconcOverall + logMDA:sex + cgconcOverall:sex+ logMDA:cgconcOverall:sex, | -185.47 |
| FW.bright ~ logMDA + cgconcOverall + sex + logMDA:cgconcOverall + logMDA:sex + cgconcOverall:sex, | -185.88 |
| **FW.bright ~ logMDA + cgconcOverall + sex,** | **-189.16** |
